# Identification of physical interactions between genomic regions by enChIP-Seq

**DOI:** 10.1101/036160

**Authors:** Toshitsugu Fujita, Miyuki Yuno, Yutaka Suzuki, Sumio Sugano, Hodaka Fujii

## Abstract

Physical interactions between genomic regions play critical roles in the regulation of genome functions, including gene expression. However, the methods for confidently detecting physical interactions between genomic regions remain limited. Here, we demonstrate the feasibility of using engineered DNA-binding molecule-mediated chromatin immunoprecipitation (enChIP) in combination with next-generation sequencing (NGS) (enChIP-Seq) to detect such interactions. In enChIP-Seq, the target genomic region is captured by an engineered DNA-binding complex, such as a CRISPR system consisting of a catalytically inactive form of Cas9 (dCas9) and a single guide RNA (sgRNA). Subsequently, the genomic regions that physically interact with the target genomic region in the captured complex are sequenced by NGS. Using enChIP-Seq, we found that the 5’HS5 locus, which regulates expression of the *β-globin* genes, interacts with multiple genomic regions upon erythroid differentiation in the human erythroleukemia cell line K562. Genes near the genomic regions inducibly associated with the 5’HS5 locus were transcriptionally up-regulated in the differentiated state, suggesting the existence of a coordinated transcription mechanism directly or indirectly mediated by physical interactions between these loci. Our data suggest that enChIP-Seq is a potentially useful tool for detecting physical interactions between genomic regions in a non-biased manner, which would facilitate elucidation of the molecular mechanisms underlying regulation of genome functions.

## Introduction

Physical interactions between genomic regions play important roles in the regulation of genome functions, including transcription and epigenetic regulation [1]. Several techniques, such as fluorescence *in situ* hybridization (FISH) [2, 3], chromosome conformation capture (3C), and 3C-derived methods [4, 5], have been used to detect such interactions. Although these techniques are widely used, they have certain limitations. The resolution of FISH is low, i.e., apparent co-localization of FISH signals does not necessarily mean that the loci in question physically interact. In addition, FISH cannot be used in a non-biased search for interacting genomic regions. In 3C and related methods, molecular interactions are maintained by crosslinking with formaldehyde prior to digestion with a restriction enzyme(s). The digested DNA is purified after ligation of the DNA ends within the same complex. Interaction between genomic loci is detected by PCR using locus-specific primers or next-generation sequencing (NGS). As with FISH, 3C-based approaches also have intrinsic drawbacks. For example, these methods require enzymatic reactions, including digestion with restriction enzyme(s) and ligation of crosslinked chromatin; the difficulty of achieving complete digestion of crosslinked chromatin can result in detection of artifactual interactions. In addition, in 3C and its derivatives, it is difficult to distinguish the products of intra-molecular and inter-molecular ligation reactions, making it difficult to ensure that the detected signals truly reflect physical interactions between different genomic regions.

An alternative approach to detecting physical interactions between genomic regions is to purify specific genomic regions engaged in molecular interactions and then analyze the genomic DNA in the purified complexes. To purify specific genomic regions, we recently developed two locus-specific chromatin immunoprécipitation (locus-specific ChIP) technologies, insertional ChIP (iChIP) [6-10] (see review [11]) and engineered DNA-binding molecule-mediated ChIP (enChIP) [12-16] (see reviews [17, 18]). enChIP consists of the following steps (Figure 1): (i) A DNA-binding molecule or complex (DB) that recognizes a target DNA sequence in a genomic region of interest is engineered. Zinc-finger proteins [19], transcription activator-like (TAL) proteins [20], and a clustered regularly interspaced short palindromic repeats (CRISPR) system [21] consisting of a catalytically inactive Cas9 (dCas9) plus a single guide RNA (sgRNA) can be used as the DB. Tag(s) and a nuclear localization signal (NLS)(s) can be fused with the engineered DB, and the fusion protein(s) can be expressed in the cells of interest. (ii) If necessary, the DB-expressing cells are stimulated and crosslinked with formaldehyde or other crosslinkers. (iii) The cells are lysed, and chromatin is fragmented by sonication or digested with nucleases. (iv) Chromatin complexes containing the engineered DB are affinity-purified by immunoprecipitation or other methods. (v) After reverse crosslinking (if necessary), DNA, RNA, proteins, or other molecules are purified and identified by various methods including NGS and mass spectrometry.

**Figure 1.**
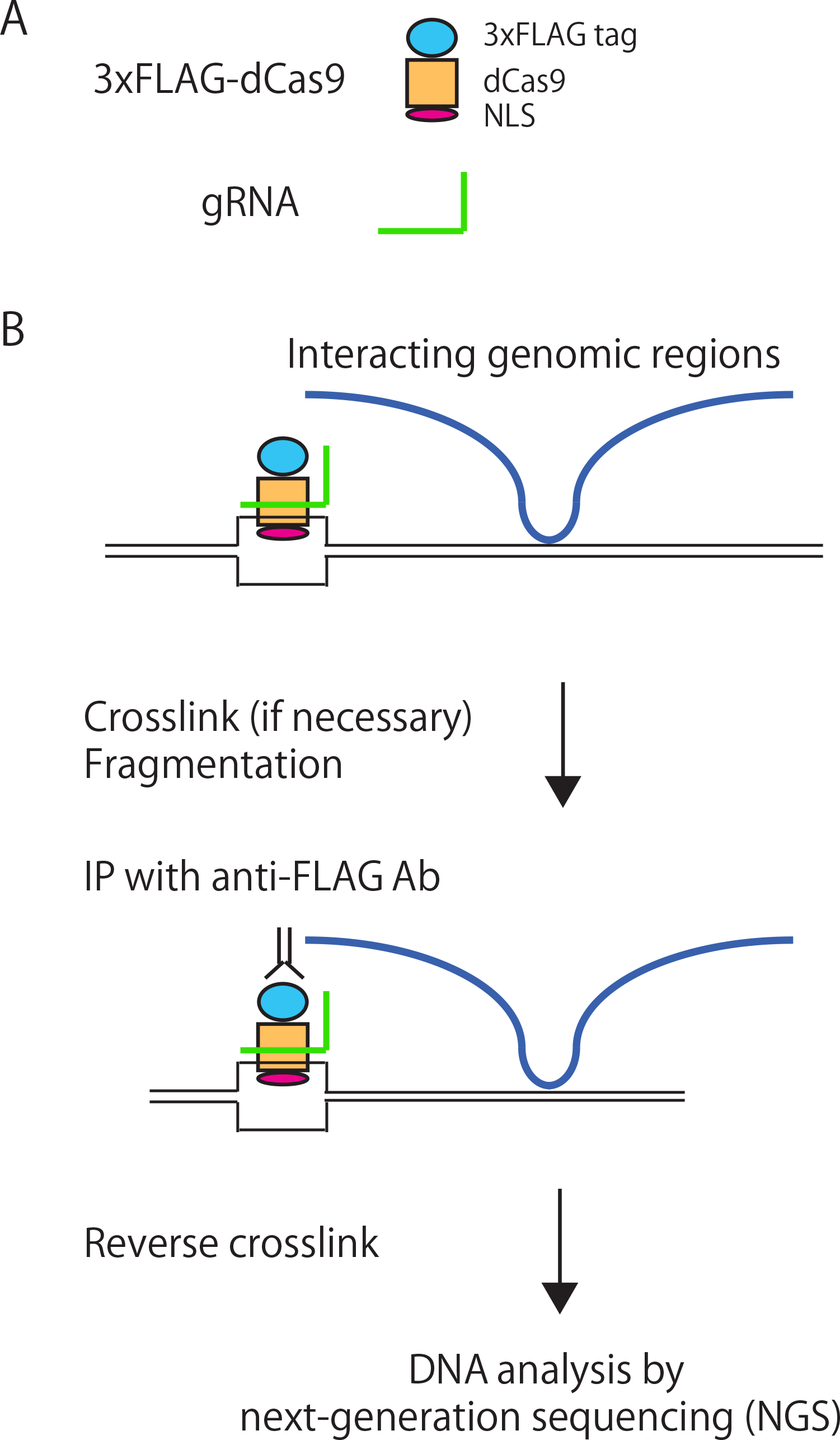
Schematic of the use of enChIP-Seq analysis to identify physical interactions between genomic regions. (**A**) The system consists of 3xFLAG-dCas9 (a fusion protein of the 3xFLAG-tag, dCas9, and a nuclear localization signal [NLS]) and a single guide RNA (sgRNA). (**B**) 3xFLAG-dCas9 and the sgRNA are expressed in the cells to be analyzed. The cells are crosslinked, if necessary, and lysed. Chromatin is purified and fragmented by sonication or other methods. Complexes containing the CRISPR complex are immunoprecipitated with anti-FLAG Ab. After reversal of crosslinking, if necessary, DNA is purified and subjected to the NGS analysis.

As a model locus in this study, we focused on 5’HS, which plays critical roles in developmentally regulated expression of the *β-globin* genes and has been extensively analyzed (Figure 2A) [22, 23]. The 5’HS2-4 regions in the 5’HS locus behave as enhancers for β-globin expression [24-26]. By contrast, 5’HS5 functions as an insulator to prevent invasion of heterochromatin into the *β-globin* genes [27]. In addition, the 5’HS5 locus interacts with the 3’HS1 locus in the 3’ region of the *globin* locus [28, 29]. Moreover, CTCF, a major component of the insulator complex, plays a critical role in insulation and formation of a chromatin loop [27, 29]. However, the molecular mechanisms underlying the functions of 5’HS5 remain incompletely understood.

**Figure 2.**
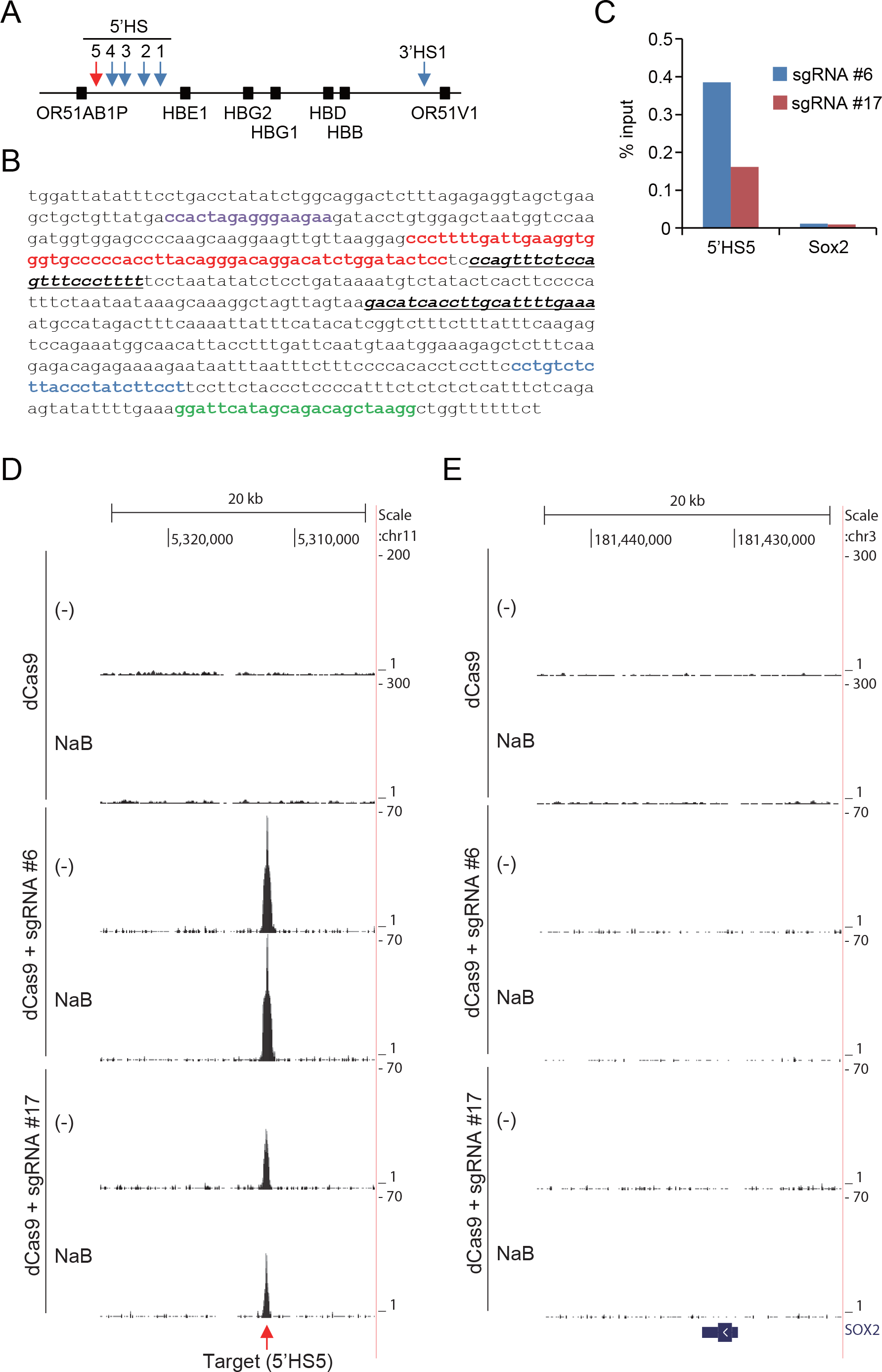
Isolation of the 5’HS5 locus by enChIP. (**A**) Schematic depiction of the *ß-globin* locus. The positions of the 5’HS and 3’HS1 loci and the *ß-globin* genes are indicated. (**B**) Positions of sgRNA target sites. Purple: CTCF-binding site [29]; red: 5’HS5 core region (NCBI Reference Sequence: NG_000007.3); blue: sgRNA target site (5’HS5 #6); green: sgRNA target site (5’HS5 #17); italic and underline: primer positions used in enChIP-real-time PCR in (**C**). (**C**) Yields of enChIP for the 5’HS5 locus. (**D** and **E**) NGS peaks at the target 5’HS5 locus (**D**) and the irrelevant *Sox2* locus (**E**). Raw ChIP-Seq read data were displayed as density plots in the UCSC Genome Browser. The vertical viewing range (y-axis shown as Scale) was set at 1-300 based on the magnitude of the noise peaks. Black vertical bars show locus positions in the human genome (hg19 assembly). The position of the *Sox2* gene is shown under the plot (**E**).

Here, we combined enChIP with NGS (enChIP-Seq) to detect genomic regions that physically interact with the 5’HS5 locus. Using enChIP-Seq, we showed that the 5’HS5 locus physically interacts with multiple genomic regions upon erythroid differentiation in the human erythroleukemia cell line K562. Thus, enChIP-Seq represents a potentially useful tool for analysis of genome functions.

## Results and Discussion

### Isolation of the 5’HS5 locus by enChIP using the CRISPR system

To purify the 5’HS5 locus by enChIP using the CRISPR system (Figures 1 and 2A) [12, 14], we generated human erythroleukemia K562-derived cells expressing 3xFLAG-dCas9 [14] and sgRNA targeting the 5’HS5 locus (Figure 2A). We designed two sgRNAs (#6 and #17) to target different sites in the 5’HS5 locus separated by 52 bp (Figure 2B). Like the parental K562 cells, the derivative cells were white, suggesting that they did not spontaneously express *globin* genes as a result of introduction of the CRISPR complex or its binding to the target locus. Crosslinked chromatin was fragmented by sonication and subjected to affinity purification with anti-FLAG antibody (Ab) (Figure 1B). Subsequently, crosslinking was reversed and DNA was purified from the isolated chromatin. As shown in Figure 2C, 0.1-0.4% of the input 5’HS5 locus was isolated, whereas the irrelevant *Sox2* locus was not enriched, suggesting that enChIP using either of the two sgRNAs (#6 and #17) could isolate the 5’HS5 locus.

### Detection of genomic regions that physically interact with the 5’HS5 locus

To identify genomic regions associated with the 5’HS5 locus on a genome-wide scale in erythroid cells under undifferentiated or differentiated conditions, K562-derived cells were mock-treated or treated with sodium butyrate (NaB) for 4 days before being crosslinked with formaldehyde and subjected to enChIP. The K562-derived cells changed from white to pink upon NaB treatment, suggesting that they had begun to express the *globin* genes, as wild-type K562 cells do in response to NaB. After isolating the 5’HS5 locus by enChIP, we subjected the purified DNA to NGS analysis. As expected, reads corresponding to the 5’HS5 locus were clearly detected in cells expressing either sgRNA #6 or #17, but not in cells expressing neither sgRNA (Figure 2D and Table 1). By contrast, no peak was detected at the irrelevant *Sox2* locus (Figure 2E).

**Table 1.** NGS information regarding the 5’HS5 target locus.

| sgRNA | Condition | Fold enrichment<br>(compared with input<br>gDNA) | Number of tags |
| --- | --- | --- | --- |
| #6 | minus (mock) | 120 | 655 |
|  | NaB | 168 | 783 |
| #17 | minus (mock) | 65 | 273 |
|  | NaB | 122 | 217 |

To identify genomic regions physically interacting with 5’HS5 upon erythroid differentiation, we analyzed the NGS peaks observed in the differentiated state. The CRISPR complex can interact with multiple genomic sites containing sequences similar to the sgRNA sequence [30-33]. In addition, in the absence of sgRNA, it is possible that dCas9 could bind non-specifically to some genomic sites *in vivo.* Therefore, the peaks identified for sgRNA #6 and #17 may include such off-target sites. To remove those off-target sites, we first eliminated peaks derived from non-specific binding of dCas9 in the absence of sgRNA from those identified for each sgRNA (Steps 1 and 2 in Figure 3). To identify peaks with confidence, we next established two criteria for choosing peaks based on NGS information from the target 5’HS5 locus: (1) tag number >5% of that of the target 5’HS5 locus, and (2) fold enrichment relative to input genomic DNA >10. As shown in Figure 3 (Step 2), 19 and 228 peaks for sgRNA #6 and #17, respectively, fulfilled these criteria. Next, to eliminate sgRNA-dependent off-target sites, we compared the peaks for sgRNA #6 and #17 and selected peaks detected in common by both sgRNA #6 and #17. These peaks were considered to represent regions engaged in bona fide physical interactions with the 5’HS5 locus (Step 3 in Figure 3). The six identified peaks could be classified into two categories: (1) peaks that were larger in the differentiated state, and (2) peaks constitutively observed in both the undifferentiated and differentiated states. The first category should contain genomic regions that inducibly associate with the 5’HS5 locus upon erythroid differentiation, whereas the second category should contain genomic regions constitutively associated with the 5’HS5 locus. To extract the peaks that grew larger specifically in the differentiated state, we selected peaks observed constitutively or in the undifferentiated state (Step 4 in Figure 3) and compared them with the six peaks extracted in Step 3 (Step 5 in Figure 3). As shown in Figure 3 (Step 5) and Table 2, the 5’HS5 site was the unique peak constitutively observed in the undifferentiated and differentiated states, whereas the five other peaks were larger specifically in the differentiated state. These included one intra-chromosomal interaction and four inter-chromosomal interactions (Table 2); the two peaks on chromosome 1 corresponding to inter-chromosomal interactions were adjacent to each other in the primary sequence. Next, we attempted to extract genomic regions that interacted with the 5’HS5 locus specifically in the undifferentiated state. However, bioinformatics analysis based on the aforementioned criteria identified no regions in this category (Figure S1).

**Table 2.** List of peak positions detected in common by both sgRNA #6 and #17.

| sgRNA #6 (NaB) |  |  |  |  |  | sgRNA #17 (NaB) |  |  |  |  |  | status |
| --- | --- | --- | --- | --- | --- | --- | --- | --- | --- | --- | --- | --- |
| chr | start | end | length | tags | fold enrichment | chr | start | end | length | tags | fold enrichment |  |
| chr1 | 247240307 | 247240894 | 588 | 68 | 17.62 | chr1 | 247240610 | 247240804 | 195 | 18 | 12.84 | NaB-specific |
| chr1 | 247240925 | 247241339 | 415 | 49 | 12.95 | chr1 | 247240971 | 247241324 | 354 | 28 | 10.33 | NaB-specific |
| chr7 | 132001863 | 132002364 | 502 | 71 | 27.24 | chr7 | 132001929 | 132002385 | 457 | 50 | 22.29 | NaB-specific |
| chr11 | 66057056 | 66057342 | 287 | 45 | 30.65 | chr11 | 66057120 | 66057297 | 178 | 13 | 14.48 | NaB-specific |
| chr15 | 64162769 | 64163183 | 415 | 51 | 29.06 | chr15 | 64162762 | 64163061 | 300 | 31 | 27.51 | NaB-specific |
| chr11 | 5311586 | 5312647 | 1062 | 783 | 168.11 | chr11 | 5311878 | 5312599 | 722 | 217 | 122.27 | constitutive |

**Figure 3.**
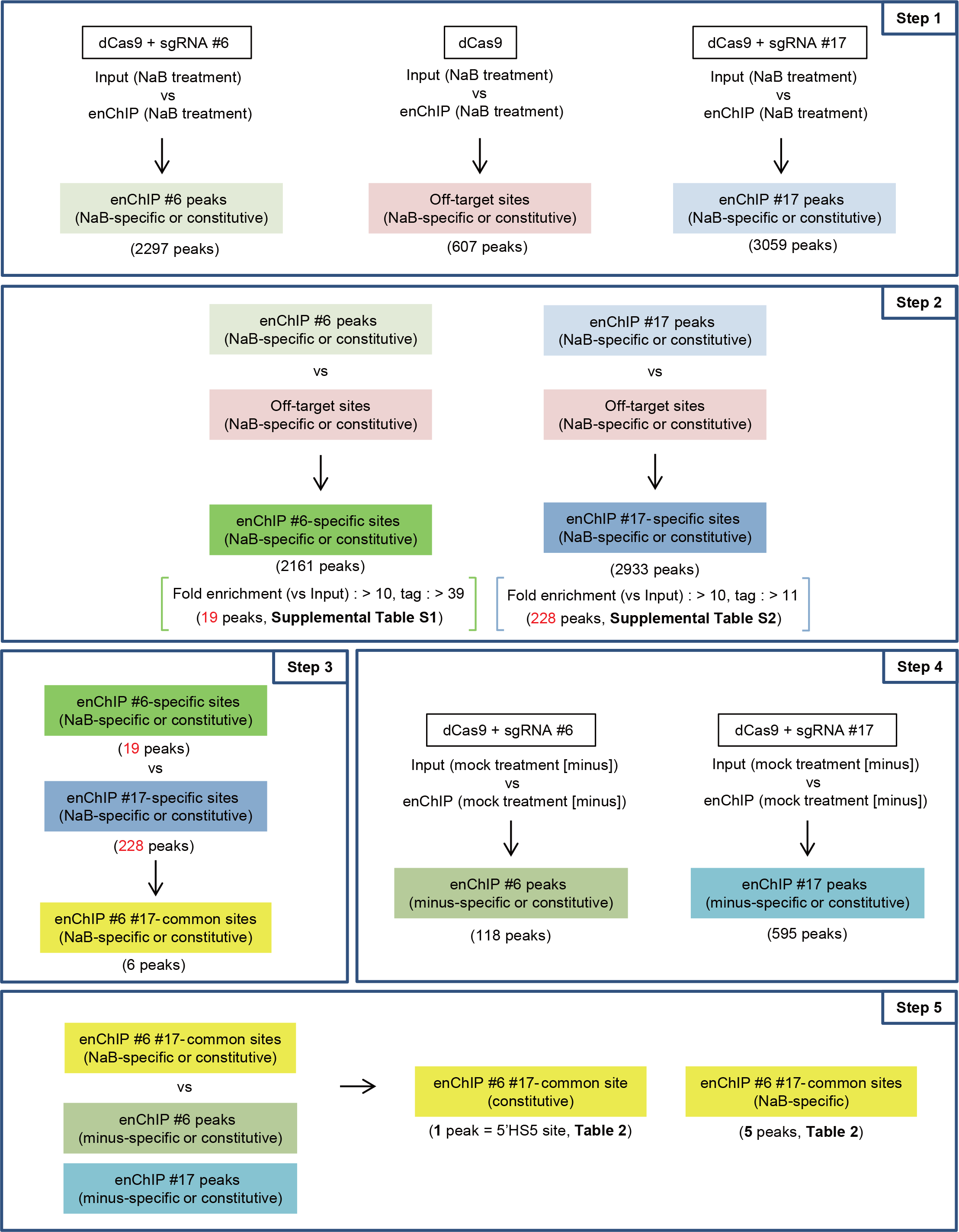
Filtering of the NGS peaks to identify genomic regions that interact with the 5’HS5 locus in the differentiated state. (**Step 1**) Extraction of enChIP-specific NGS peaks by comparison with enChIP and input peaks detected in the differentiated state, using Model-based Analysis of ChIP-Seq (MACS) (http://liulab.dfci.harvard.edu/MACS/). The extracted enChIP-specific peaks include NaB-specific and constitutively detected peaks. (**Step 2**) Elimination of peaks derived from non-specific binding of dCas9. After the elimination step, peaks fulfilling the defined criteria (19 for sgRNA #6 and 228 for sgRNA #17) were analyzed in step 3. (**Step 3**) Identification of peaks detected in common by both sgRNA #6 and #17. Genomic regions corresponding to the identified peaks were considered to physically interact with the 5’HS5 locus. (**Step 4**) Extraction of enChIP-specific NGS peaks by comparison with enChIP and input peaks detected in the undifferentiated state, using MACS. The extracted enChIP-specific peaks include peaks specifically detected in undifferentiated cells and constitutively detected peaks. (**Step 5**) Identification of genomic regions that interact with the 5’HS5 locus in a NaB-specific manner.

We visualized some of the identified peaks in the UCSC Genome Browser (Figure 4). The peaks were clearly visible for both sgRNA #6 and #17. The adjacent NaB-specific peaks in chromosome 1 were both located in the first intron of the *ZFN670* and *ZFN670-695* genes. The other NaB-specific peaks were located in the vicinity of the *MIR422A* gene in chromosome 15 and between the *TMEM151A* and *YIF1A* genes in chromosome 11.

**Figure 4.**
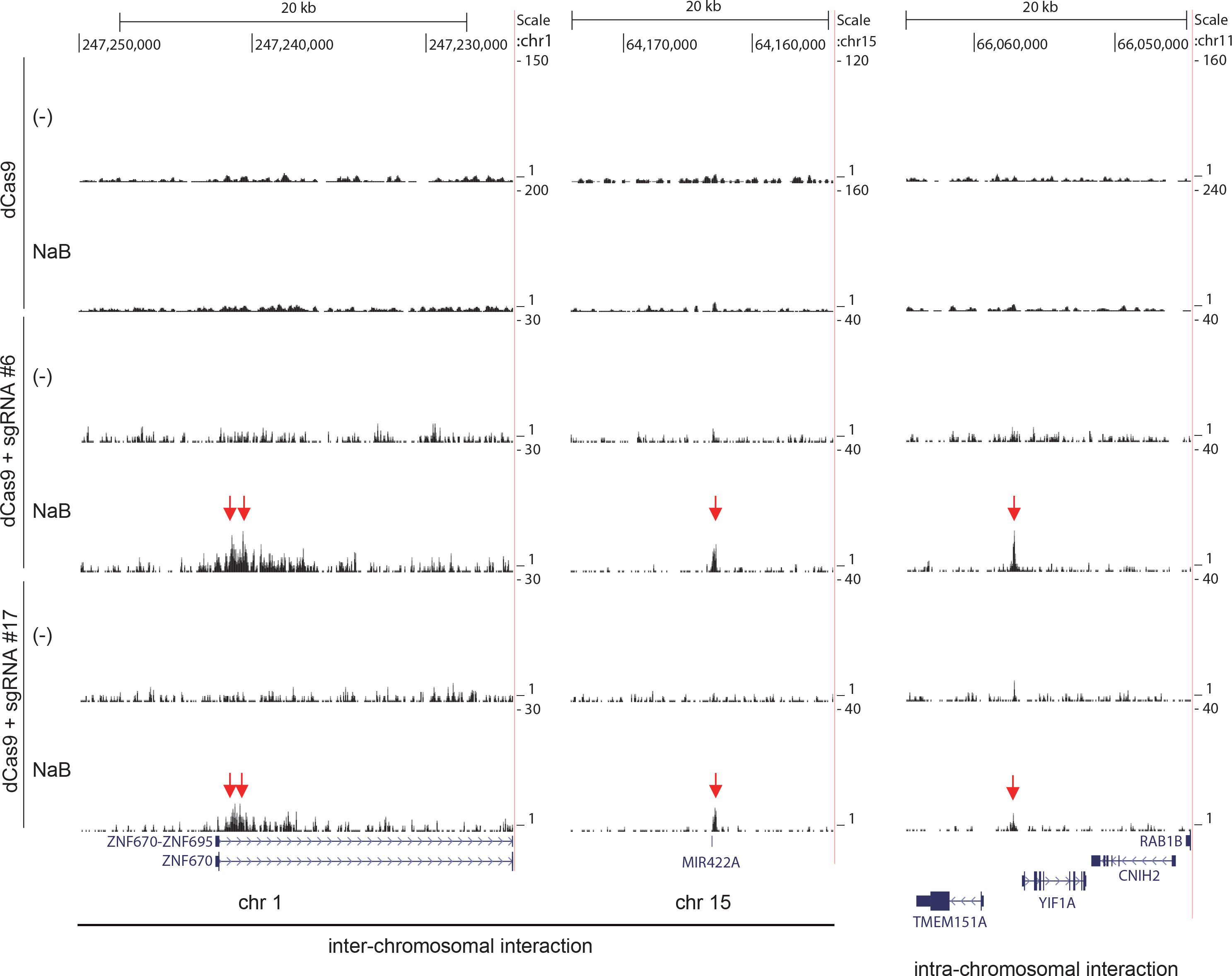
Identification of genomic regions that physically interact with the 5’HS5 locus by enChIP-Seq. (**Step 1**) Raw ChIP-Seq read data were displayed as density plots in the UCSC Genome Browser. The vertical viewing range (y-axis shown as Scale) was set at 1-160 based on the noise peaks. Black vertical bars reveal locus positions in the human genome (hg19 assembly). Positions of genes are shown under the plots.

To confirm the interactions identified by enChIP-Seq, we used the ligation-mediated approach used in 3C-based assays (Figure S2). In this approach, cells are subjected to crosslinking, and then chromatin is randomly fragmented by sonication. After proximity ligation of genomic DNA, the junction between the target locus and a potential interacting site is amplified by PCR. Subsequently, a part of the amplified region in the potential interacting locus is detected by a second PCR. Amplification of a region in the second PCR suggests that the potential interacting locus is physically proximal to the target locus. When we used this assay to examine the amplified the *ZNF670/ZFN670-695* locus in a NaB-specific manner only when the proximal ligation step was performed (Figure 5). This observation is consistent with the enChIP-Seq result (Figure 4) showing that the 5’HS5 locus interacts with the *ZNF670/ZFN670-695* locus in the differentiated state in K562 cells. Thus, we were able to confirm the chromosomal interaction identified by enChIP-Seq by another independent method, suggesting that it is feasible to use enChIP-Seq to perform non-biased identification of physical interactions between genomic regions.

**Figure 5.**
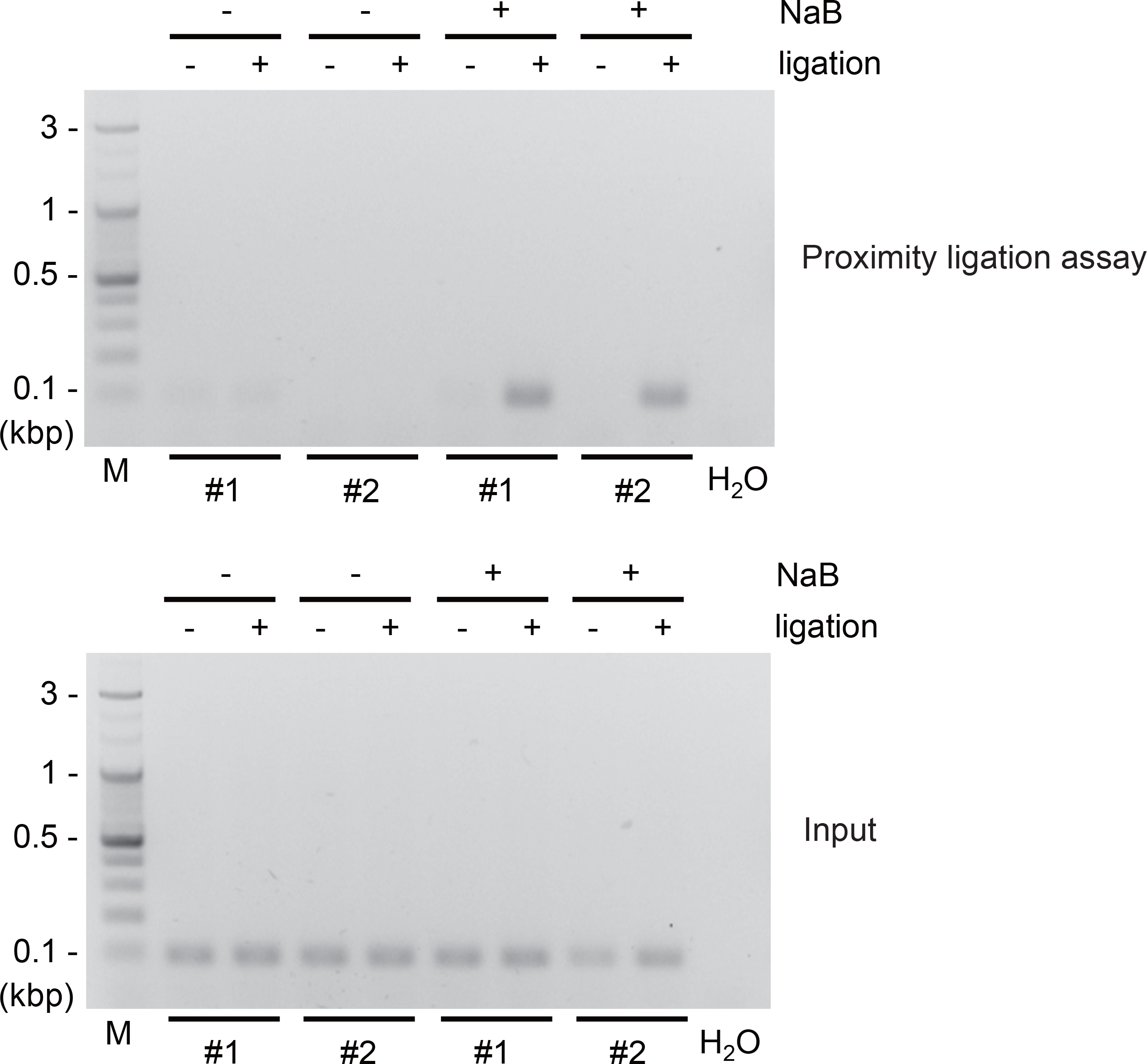
Confirmation of enChIP-Seq results by proximity ligation assay. (**Upper**) Results of PCR. K562 cells cultured in the presence or absence of NaB were crosslinked, and the chromatin was extracted, fragmented, and subjected to the proximity ligation assay (Figure S2) to confirm interaction between the 5’HS5 and *ZNF670/ZFN670-695* loci. A region of the *ZNF670/ZFN670-695* locus was detected by the second PCR. (**Lower**) The result of PCR with DNA used for the first PCR and the primer set used for the second PCR. This result shows that the reactions were run with comparable amounts of input DNA. Two representatives of each sample (±NaB treatment) are shown.

Transcription of genes near the 5’HS5-interacting genomic regions identified by enChIP-Seq could be directly or indirectly regulated by the induced association with the 5’HS locus. Such regulation could involve the 5’HS2-4 regions, which function as enhancers [24-26]. Therefore, we investigated whether mRNA levels of the genes in the vicinity (±10 kbp) of the 5’HS5-interacting genomic regions changed after NaB treatment. As shown in Figure 6, mRNA levels of the *ZNF670, MIR422A*, and *CNIH2* genes were clearly up-regulated in the NaB-induced differentiated state, suggesting that the identified chromosomal interactions are involved in transcriptional regulation of these genes. At this time, it is not clear whether these gene products play any roles in erythroid development. Future studies should attempt to elucidate how the 5’HS locus regulates transcription of these genes. It is possible that the enhancer function of the 5’HS locus directly activates transcription of these genes via interactions with their promoters under differentiated conditions. Alternatively, these loci may be incorporated into the “transcription factory” [34] upon erythroid differentiation, independent of the enhancer function of the 5’HS locus.

**Figure 6.**
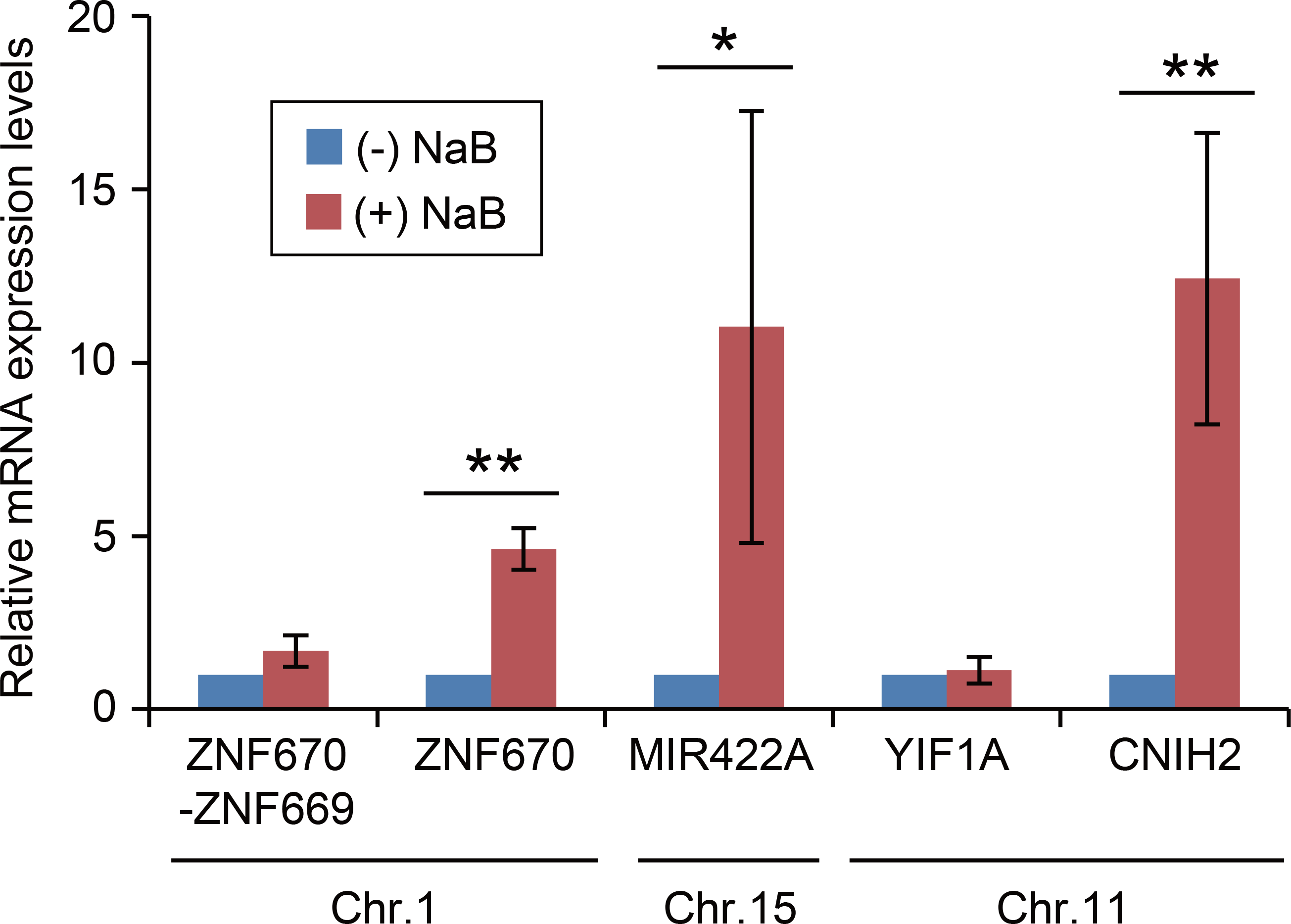
Confirmation of enChIP-Seq results by proximity ligation assay. (**Upper**) NaB-induced expression of genes in the vicinity of genomic regions that interact with the 5’HS5 locus. Total RNA was used for quantitative RT-PCR analysis. Expression levels of the indicated genes were normalized to those of *GAPDH*, and the levels of mRNA in the absence of NaB were defined as 1 (mean ± SD, n = 3). N.D.: not detected, *: t-test p value <0.05, **: t-test p value <0.01.

### Lack of interaction between the 5′HS5 and *β-globin* loci

Studies using 3C and derived methods suggested that the 5’HS5 and *β-globin* loci interact [28, 29]. Therefore, we sought to detect a physical interaction between these loci by enChIP-Seq. As shown in Figure 3, bioinformatics analysis based on the criteria described above (fold enrichment: >10, tag number: >5% of that of the target positions) did not extract genomic regions around the *β-globin* locus. In fact, the peak images did not indicate any physical interactions between 5’HS5 and the *β-globin* locus or 3’HS1 (Figure S3). Importantly, in this regard, it is not likely that bound dCas9/sgRNA complexes abrogate the formation of chromosomal loops between 5’HS5 and the *β-globin* or 3’HS1 locus, because the target positions of the sgRNA do not overlap with the CTCF-binding site and the 5’HS5 core region (Figure 2), and the cells maintained their capability to differentiate in response to NaB.

Several phenomena might explain the discrepancy between the results of this study and those of 3C-derived methods regarding the interaction of 5’HS5 with the *β-globin* locus. First, 3C-derived methods might be much more sensitive than enChIP-Seq. Specifically, because 3C-derived methods use PCR amplification to detect ligated regions consisting of different genomic regions, they may be able to detect transient or weak interactions that did not pass the criteria we used in this study. Second, the fragmentation of chromatin by sonication in enChIP-Seq may be too harsh to retain weak chromosomal interactions. By contrast, 3C-derived methods employ restriction digestion, which is much milder than sonication, to fragment chromatin. Third, physical interactions between genomic regions are likely to be regulated in a cell cycle-dependent manner, and it is possible that interactions between these regions may occur only in a certain phase of a cell cycle, making it difficult to detect by enChIP-Seq. Alternatively, our results raise the possibility that the ‘interactions’ detected by 3C and its derivatives reflect accessibility of the loci to the nucleases and ligases employed in these techniques, but do not necessarily reflect physical interactions between genomic regions. In fact, discrepancies between results of 3C or its derivatives and those of FISH have been suggested [35]. This possibility highlights the importance of confirming chromosomal interactions by independent methods.

### Managing potential contamination of off-target sites

In this study, we identified physical interactions between genomic regions based on signals detected by enChIP-Seq. dCas9 can bind to multiple sites containing sequences similar to the sgRNA sequence [30, 31]. To eliminate potential contamination of our findings by off-target sites, we propose several strategies:

(1) Carefully examine the sequences of the detected peaks and remove those containing sequences similar to the target sequence.

(2) Use different conditions or cell types. Signals specifically detected in one condition or cell type should reflect true physical interactions between genomic regions.

(3) Use multiple different sgRNAs. Because different sgRNA are unlikely to engage in off-target binding at the same genomic regions, signals observed in common using different sgRNAs should reflect true physical interactions between genomic regions. In addition, cells expressing dCas9 without sgRNA should be used as a negative control to eliminate off-target sites associated with dCas9 in the absence of sgRNA.

(4) Use a sequential purification scheme. Cas9 orthologs derived from different bacterial species recognize distinct proto-spacer adjacent motif (PAM) sequences and can be used for genome editing and gene regulation [36]. Tagging of a given locus with dCas9s derived from different lineages and bearing distinct tags would make it feasible to sequentially purify the locus, minimizing contamination with off-target sites.

Using these techniques, we believe that we can effectively manage potential contamination of dCas9 off-target sites in enChIP analyses.

## Conclusions

In this study, we used enChIP-Seq analysis to detect physical interactions between genomic regions. In K562-derived cells, the 5’HS5 locus physically interacted with multiple regions in the genome (Figure 4, Table 2). These interactions were induced by erythroid differentiation in response to NaB treatment (Figure 4, Table 2). Transcription of genes around the interacting genomic region was up-regulated in the differentiated state (Figure 6), suggesting a direct or indirect involvement of 5’HS enhancer activity in transcription of genes proximal to the interacting sites. Our results suggest that enChIP-Seq represents a potentially useful tool for performing non-biased searches for physical interactions between genomic regions, which would facilitate elucidation of the molecular mechanisms underlying regulation of genome functions.

## Materials and Methods

### Plasmids

3xFLAG-dCas9/pMXs-puro (Addgene #51240) was described previously [14]. To construct vectors for expression of sgRNAs, two oligos for each sgRNA were annealed and extended using Phusion polymerase (New England Biolabs) to make 100 bp double-stranded DNA fragments, as described previously [12]. The nucleotide sequences were as follows: hHS5 #6, 5’-TTTCTTGGCTTTATATATCTTGTGGAAAGGACGAAACACCGGATTCATAGCAGACA GCTA-3’ and 5’-GACTAGCCTTATTTTAACTTGCTATTTCTAGCTCTAAAACTAGCTGTCTGCTATGAA TCC-3’; hHS5 #17, 5’-TTTCTTGGCTTTATATATCTTGTGGAAAGGACGAAACACCGGGAAGATAGGGTAA GAGAC-3’ and 5’-GACTAGCCTTATTTTAACTTGCTATTTCTAGCTCTAAAACGTCTCTTACCCTATCTT CCC-3’. Fragments were purified following agarose gel electrophoresis and subjected to Gibson assembly (New England Biolabs) with the linearized sgRNA cloning vector (Addgene #41824), a gift from George Church [37], to yield sgRNA-hHS5 #6 and sgRNA-hHS5 #17. The gBlocks were excised with *XhoI* and *HindIII* and cloned into *XhoI/HindIII-cleaved* pSIR vector to generate self-inactivating retroviral vectors for sgRNAs, as described previously [14].

### Cell culture

K562-derived cells [14] were maintained in RPMI (Wako) supplemented with 10% fetal calf serum (FCS).

### Establishment of cells stably expressing 3xFLAG-dCas9 and sgRNA

Establishment of K562-derived cells expressing 3xFLAG-dCas9 was described previously [14]. To establish cells expressing both 3xFLAG-dCas9 and sgRNAs targeting the 5’HS5 locus, 2 μg of sgRNA-hHS5 #6/pSIR or sgRNA-hHS5 #17/pSIR was transfected along with 2 μg of pPAM3 into 1 × 10^6^ 293T cells [38]. Two days after transfection, K562-derived cells expressing 3xFLAG-dCas9 were infected with the supernatant (5 ml) of 293T cells containing the virus particles. K562-derived cells expressing both 3xFLAG-dCas9 and sgRNA-hHS5 #6 or sgRNA-hHS5 #17 were selected in RPMI medium containing 10% FCS, puromycin (0.5 μg/ml), and G418 (0.8 mg/ml).

### Induction of differentiation of K562-derived cells

To induce erythroid differentiation of the K562-derived cells, cells were incubated in the presence of 1 mM NaB for 4 days.

### enChIP-real-time PCR

enChIP-real-time PCR was performed as previously described [12], except that ChIP DNA Clean & Concentrator (Zymo Research) was used for purification of DNA. Primers used in the analysis are shown in Table S3.

### enChIP-Seq and bioinformatics analysis

Undifferentiated or differentiated K562-derived cells (2 × 10^7^ each) expressing 3xFLAG-dCas9 and sgRNAs were subjected to the enChIP procedure as described previously [12], except that ChIP DNA Clean & Concentrator was used for purification of DNA. NGS and data analysis were performed at the University of Tokyo as described previously [39, 40]. Additional data analysis for Step 2 in Figure 3 was performed at Hokkaido System Science Co., Ltd. Images of NGS peaks were generated using the UCSC Genome Browser (https://genome.ucsc.edu/cgi-bin/hgGateway).

### Proximity ligation assay to confirm interactions between genomic regions

K562 cells (1 × 10^7^) were fixed with 1% formaldehyde at 37°C for 5 min. The chromatin fraction was extracted and fragmented by sonication (fragment length, 2 kb on average) as described previously [41], except for the use of 800 μl of TE buffer (10 mM Tris pH 8.0, 1 mM EDTA) and a UD-201 ultrasonic disruptor (TOMY SEIKO). Sonicated chromatin (34 μl was treated with the End-It DNA End-Repair kit (Epicentre) in a 50 μl reaction mixture at room temperature for 45 min. After heating at 70°C for 10 min, reaction mixture (23.5 μl was incubated in the presence or absence of T4 DNA ligase (Roche) at room temperature for 2 h. After reverse crosslinking at 65°C followed by RNase A and Proteinase K treatment, DNA was purified using ChIP DNA Clean & Concentrator. The purified DNA was used as a template for the first PCR with KOD FX (Toyobo) and a primer set including one primer containing an *I-SceI* site that was biotinylated at the 5’ end (Table S3). PCR conditions were as follows: denaturing at 94°C for 2 min; 30 cycles of 98°C for 10 sec, 60°C for 30 sec, and 68°C for 6 min. The reaction mixture (15 μl) was mixed with 15 μl of Dynabeads M-280 Streptavidin (Thermo Fisher Scientific) and 500 μl of RIPA buffer (50 mM Tris [pH 7.5], 150 mM NaCl, 1 mM EDTA, 0.5% sodium deoxycholate, 0.1% SDS, 1% IGEPAL-CA630) at 4°C for 1 h. After three washes with RIPA buffer and one wash with 1 × NEBuffer 2 (New England Biolabs), the Dynabeads were treated with *I-Sce I* at 37°C for 2 h. The supernatant was collected, incubated at 65°C for 20 min, and used for the second PCR with AmpliTaq Gold 360 Master Mix (Applied Biosystems). PCR conditions were as follows: denaturing at 95°C for 10 min; 27 cycles of 95°C for 30 sec, 60°C for 30 sec, and 72°C for 1 min. Primers used in the analysis are shown in Table S3.

### RNA extraction and quantitative RT-PCR

Total RNA was extracted from mock- or NaB-treated K562 cells and used for quantitative RT-PCR as previously described [42]. Primers used in the analysis are shown in Table S3.

## Acknowledgments

We thank G.M. Church for providing a plasmid (Addgene plasmid #41824), F. Kitaura for technical assistance, and T. Kikuchi, H. Horiuchi, and M. Tosaka for NGS analysis.

---

**Table S1.**
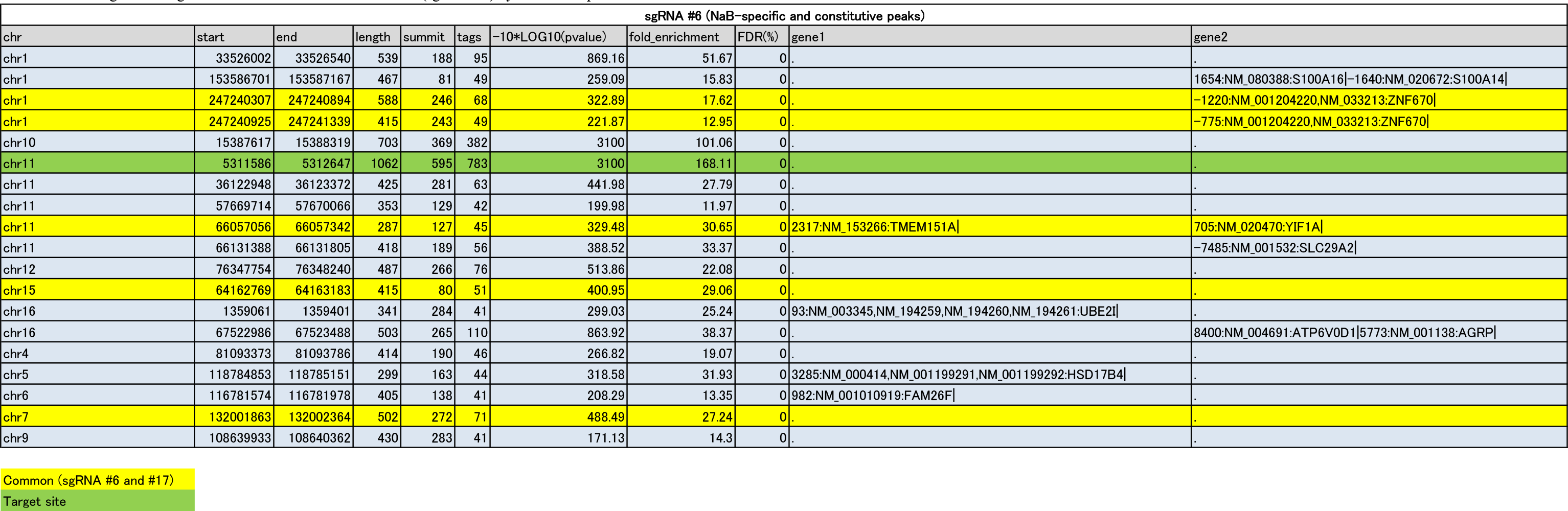
List of genomic regions detected in the differentiated state (sgRNA #6) by enChIP-Seq. (.xlsx)

**Table S2.**
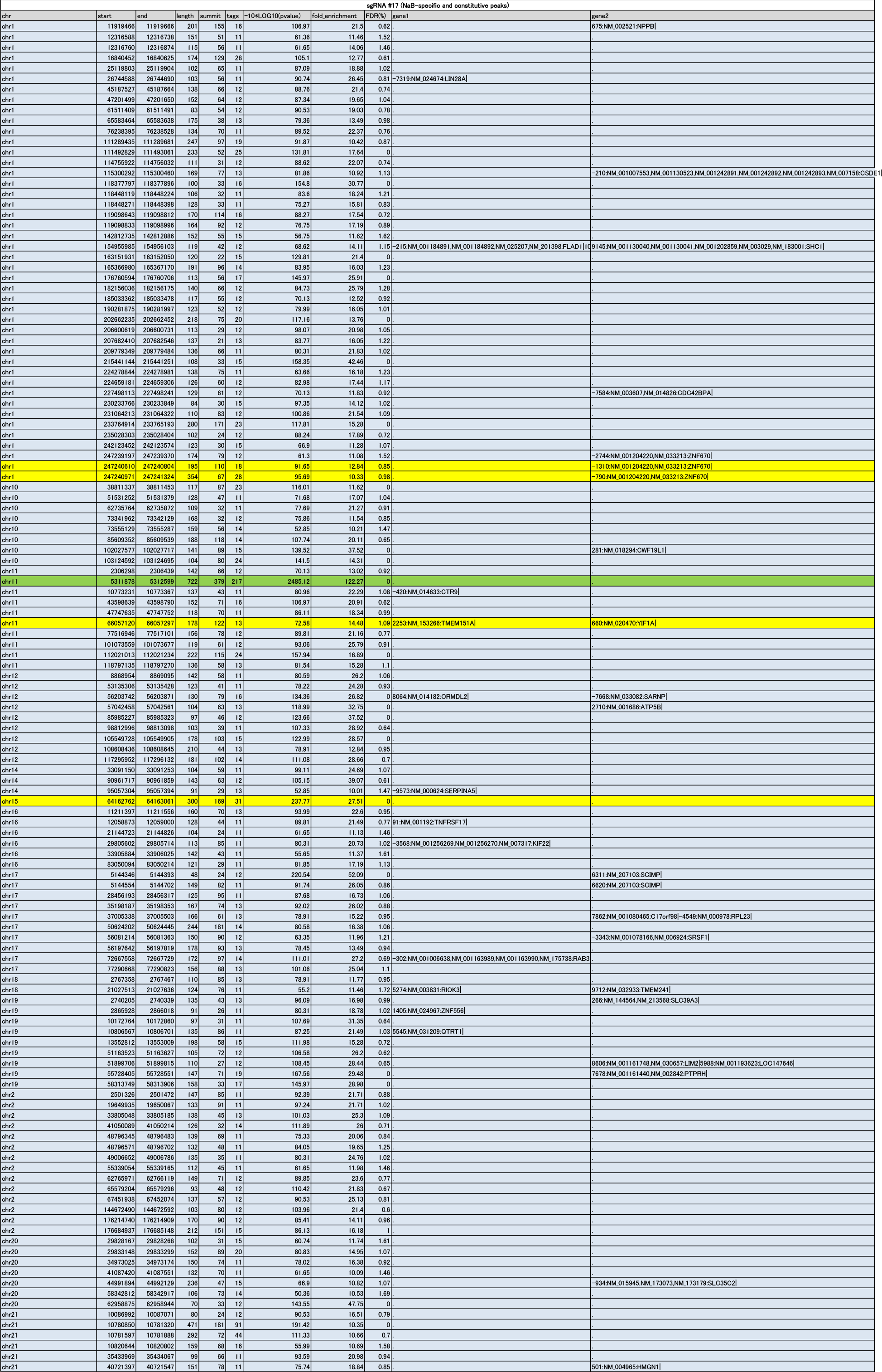
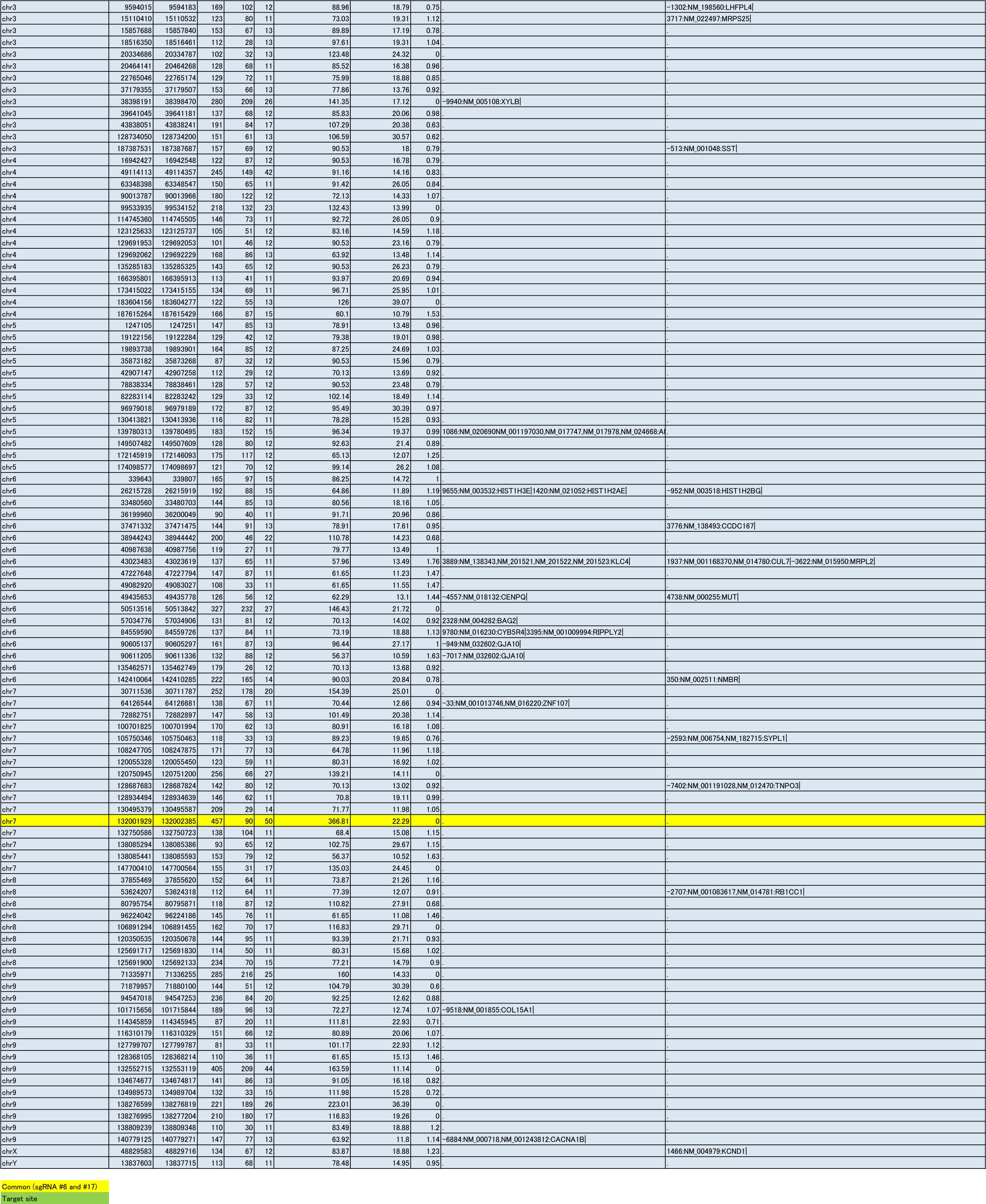
List of genomic regions detected in the differentiated state (sgRNA #17) by enChIP-Seq. (.xlsx)

**Table S3.**
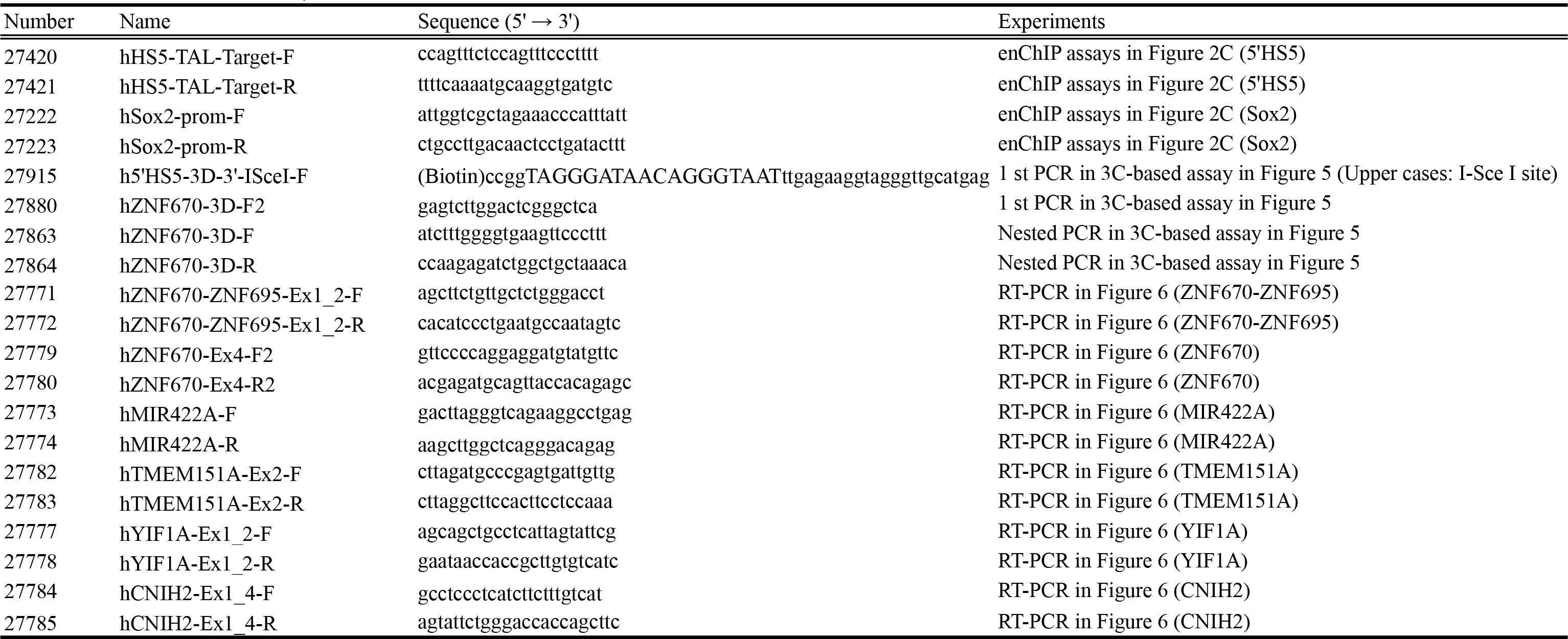
Primers used in this study. (.xlsx)

**Figure S1.**
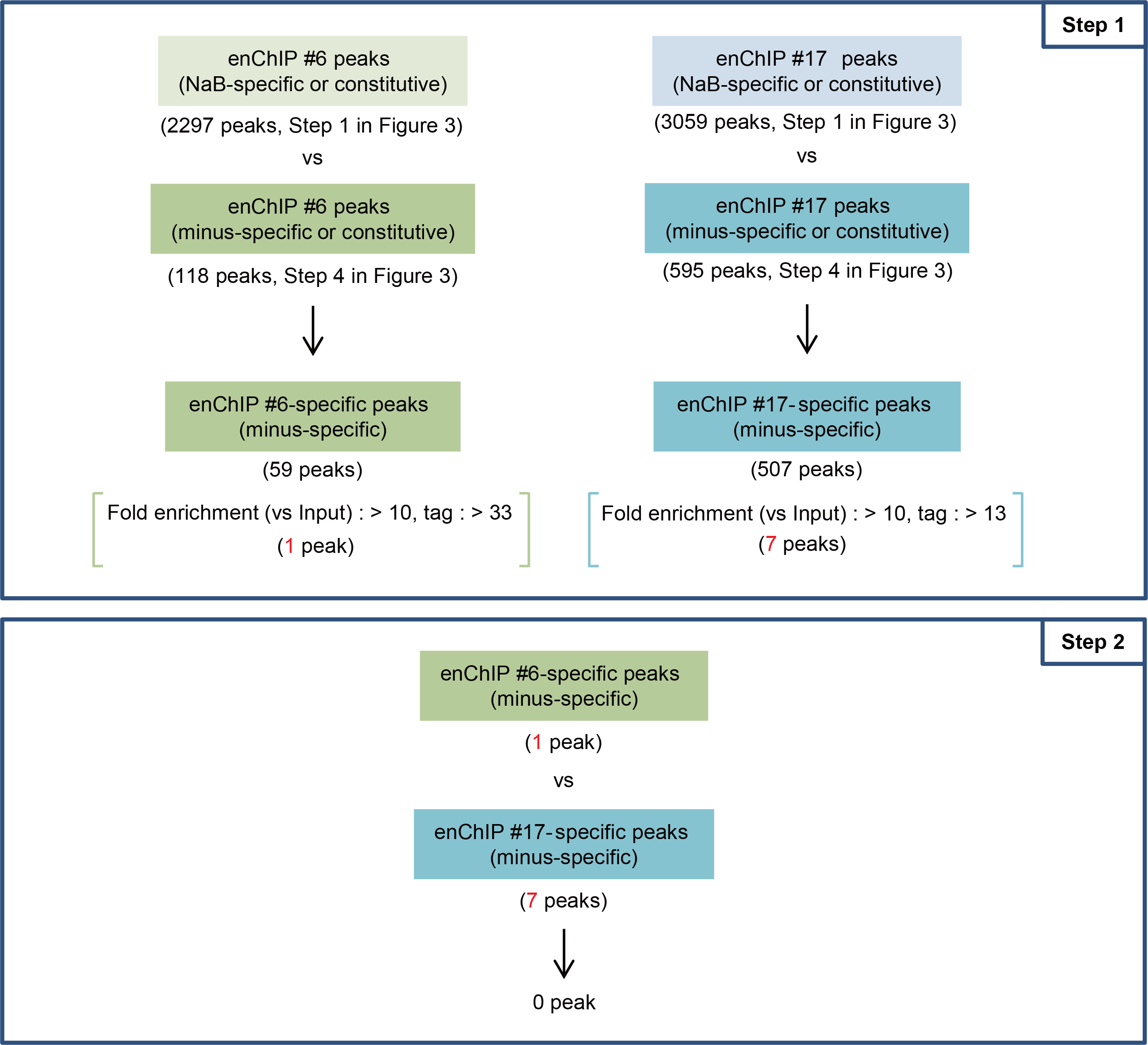
Filtering of NGS peaks to identify genomic regions that interact with the 5’HS5 locus in the undifferentiated state.(**Step 1**. Extraction of genomic regions that interact with the 5’HS5 locus specifically in the undifferentiated state. After the extraction step, regions fulfilling the defined criteria (one for sgRNA #6 and seven for sgRNA #17) were analyzed in step 2. (**Step 2**) Identification of peaks detected in common using sgRNA #6 and #17.

**Figure S2.**
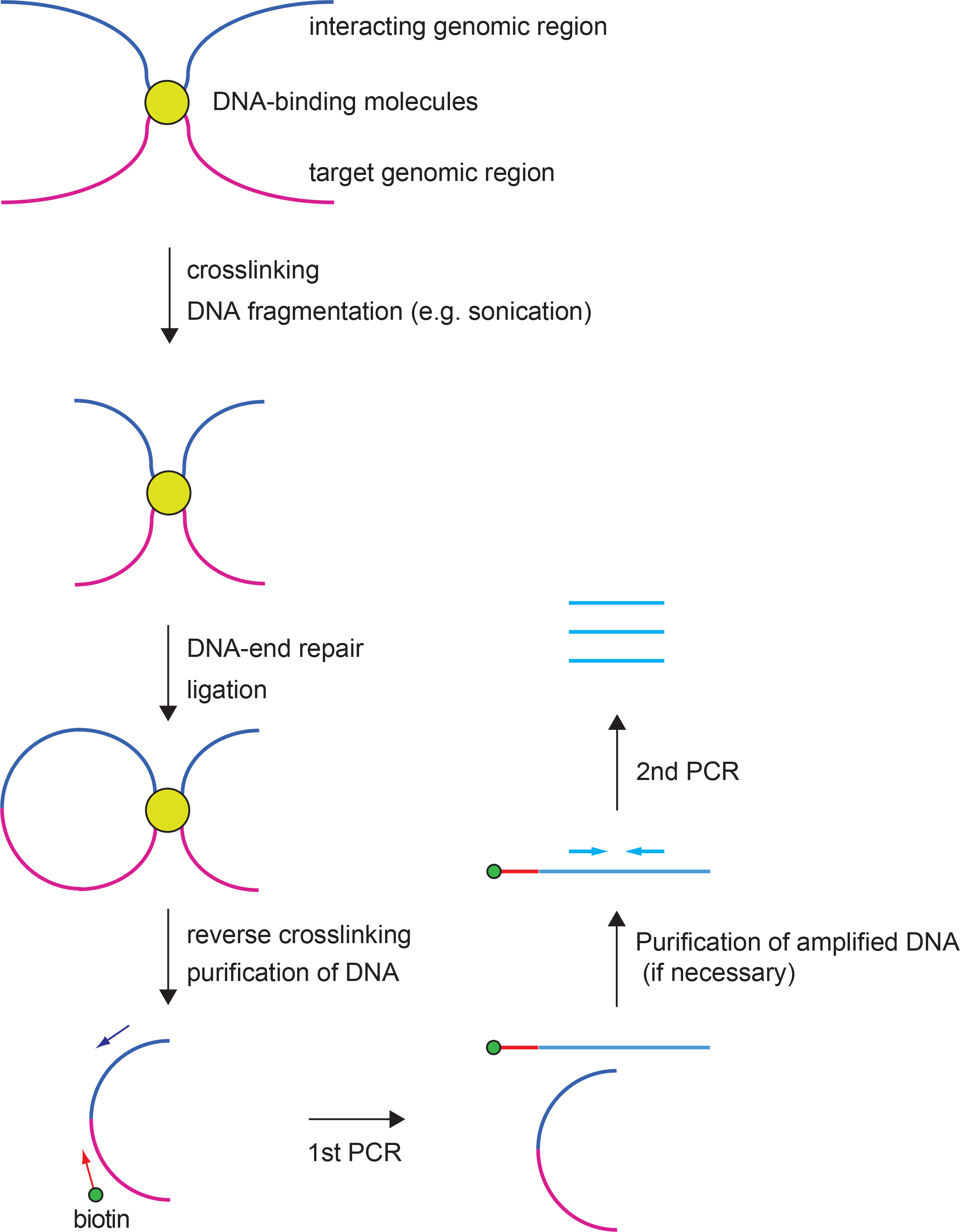
Scheme for proximity ligation assay to confirm interactions between genomic regions. Cells are crosslinked and lysed, and genomic DNA is randomly fragmented by sonication or other methods. After repair of DNA ends, proximal ligation, and reversal of crosslinking, the junction between the target locus and its potential interacting region is amplified by PCR. If a biotinylated primer is used, the amplicon can be purified using streptavidin. Subsequently, a part of the amplified region in the potential interacting locus is detected by the second PCR. Amplification of the region in the second PCR suggests that the potential interacting locus is physically proximal to the target locus.

**Figure S3.**
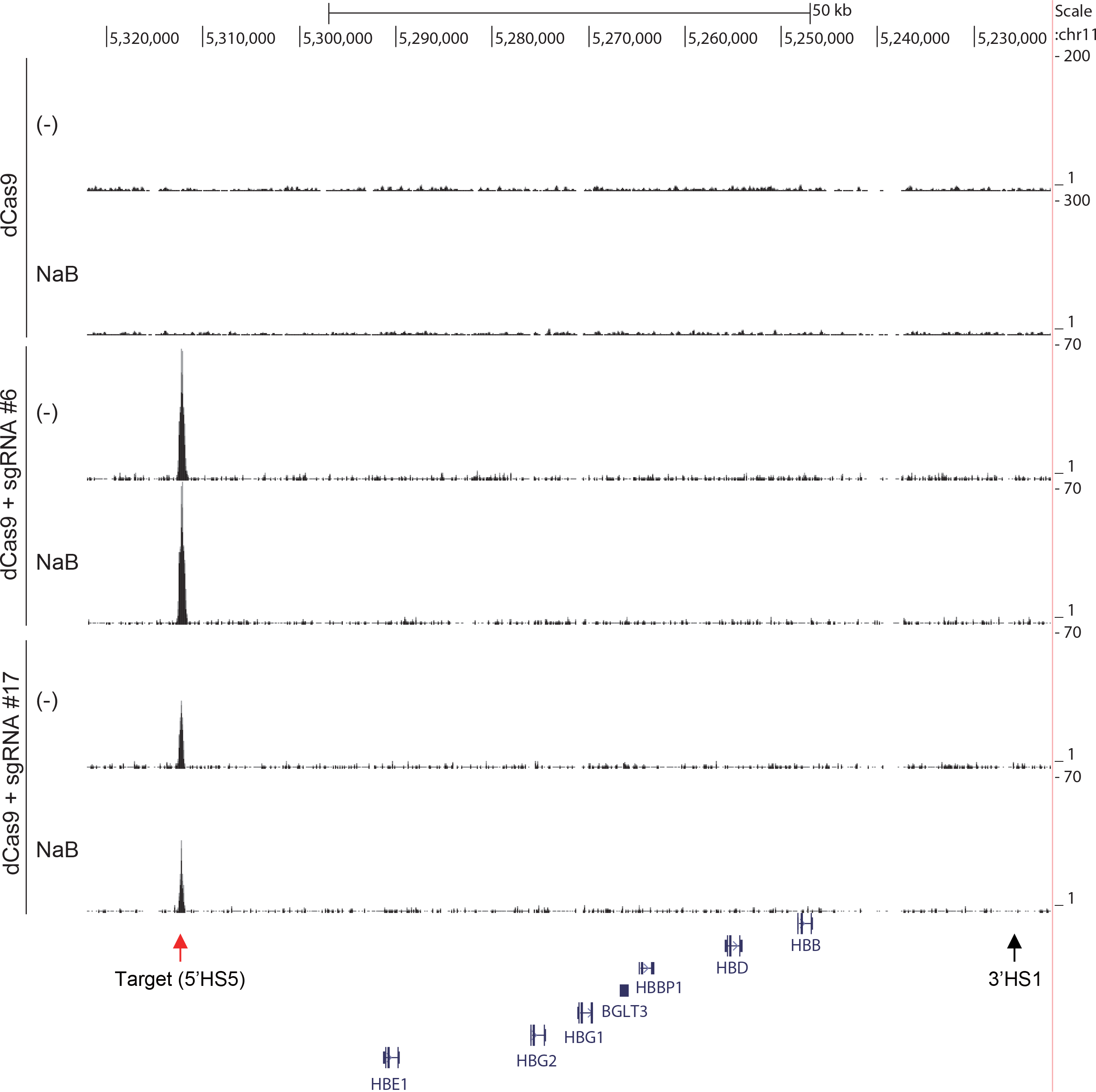
NGS peak images in the *β-globin* locus. Although a peak at 5’HS5 was clearly observed, no peak indicated a physical interaction between 5’HS5 and other regions in the *β-globin* locus. Raw ChIP-Seq read data were displayed as density plots in the UCSC Genome Browser. The vertical viewing range (y-axis shown as Scale) was set at 1-300, based on the magnitude of the noise peaks. Black vertical bars show locus positions in the human genome (hg19 assembly). Positions of genes, including the *β-globin* genes, are shown under the plots.

## References

1. Williams A, Spilianakis CG, Flavell RA. Interchromosomal association and gene regulation in trans. Trends Genet. 2010 26: 188–97.

2. Fisher AG, Merkenshlager M. Gene silencing, cell fate and nuclear organization. Curr Opin Genet Dev. 2002; 12(2): 193–7.

3. Fraser P, Bickmore W. Nuclear organization of the genome and the potential for gene regulation. Nature. 2007; 447 (7143): 413–7.

4. Dekker J, Rippe K, Dekker M, Kleckner N. Capturing chromosome conformation. Science. 2002; 295 (5558): 1306–11.

5. de Wit E, de Laat W. A decade of 3C technologies: insights into nuclear organization. Genes Dev. 2012; 26(1): 11–24.

6. Hoshino A, Fujii H. Insertional chromatin immunoprecipitation: a method for isolating specific genomic regions. J Biosci Bioeng. 2009; 108(5): 446–9. doi: 10.1016/jjbiosc.2009.05.005.

7. Fujita T, Fujii H. Direct idenification of insulator components by insertional chromatin immunoprecipitation. PLoS One. 2011; 6(10): e26109. doi: 10.1371/journal.pone.0026109.

8. Fujita T, Fujii H. Efficient isolation of specific genomic regions by insertional chromatin immunoprecipitation (iChIP) with a second-generation tagged LexA DNA-binding domain. Adv Biosci Biotechnol. 2012; 3(5): 626–9. doi: 10.4236/abb.2012.35081.

9. Fujita T, Fujii H. Efficient isolation of specific genomic regions retaining molecular interactions by the iChIP system using recombinant exogenous DNA-binding proteins. BMC Mol Biol. 2014; 15: 26. doi: 10.1186/s12867-014-0026-0.

10. Fujita T, Kitaura F, Fujii H. A critical role of the Thy28-MYH9 axis in B cell-specific expression of the *Pax5* gene in chicken B cells. PLoS One. 2015; 10: e0116579. doi: 10.1371/journal.pone.0116579.

11. Fujita T, Fujii H. Biochemical analysis of genome functions using locus-specific chromatin immunoprecipitation technologies. Gene Regul Syst Bio. 2015; in press.

12. Fujita T, Fujii H. Efficient isolation of specific genomic regions and identification of associated proteins by engineered DNA-binding molecule-mediated chromatin immunoprecipitation (enChIP) using CRISPR. Biochem Biophys Res Commun. 2013; 439: 132–6. doi: 10.1016/j.bbrc.2013.08.013.

13. Fujita T, Asano Y, Ohtsuka J, Takada Y, Saito K, Ohki R, et al. Identification of telomere-associated molecules by engineered DNA-binding molecule-mediated chromatin immunoprecipitation (enChIP). Sci Rep. 2013; 3: 3171. doi: 10.1038/srep03171.

14. Fujita T, Fujii H. Identification of proteins associated with an IF_γ_-responsive promoter by a retroviral expression system for enChIP using CRISPR. PLoS One. 2014; 9(7): e103084. doi: 10.1371/journal.pone.0103084.

15. Fujita T, Yuno M, Okuzaki D, Ohki R, Fujii H. Identification of non-coding RNAs associated with telomeres using a combination of enChIP and RNA sequencing. PLoS One. 2015; 10: e0123387. doi: 10.1371/journal.pone.0123387.

16. Fujita T, Yuno M, Fujii H. Efficient sequence-specific isolation of DNA fragments and chromatin by in vitro enChIP technology using recombinant CRISPR ribonucleoproteins. Genes Cells. 2016; in press.

17. Fujii H, Fujita T. Isolation of specific genomic regions and identification of their associated molecules by engineered DNA-binding molecule-mediated chromatin immunoprecipitation (enChIP) using the CRISPR system and TAL proteins. Int J Mol Sci. 2015 16: 21802–12.

18. Fujita T, Fujii H. Applications of engineered DNA-binding molecules such as TAL proteins and the CRISPR/Cas system in biology research. Int J Mol Sci. 2015 16: 23143–64.

19. Pabo CO, Peisach E, Grant RA. Design and selection of novel Cys2his2 zinc finger poroteins. Annu Rev Biochem. 2001 70: 313–40.

20. Bogdanove AJ, Voytas DF. TAL effectors: customizable proteins for DNA targeting. Science. 2011; 333 (6051): 1843–6.

21. Qi LS, Larson MH, Gilbert LA, Doudna JA, Weissman JS, Arkin AP, et al. Repurposing CRISPR as an RNA-guided platform for sequence-specific control of gene expression. Cell. 2013; 152: 1173–83. doi: 10.1016/j.cell.2013.02.022.

22. Ghirlando R, Giles K, Gowher H, Xiao T, Xu Z, Yao H, et al. Chromatin domains, insulators, and the regulation of gene expression. Biochim Biophys Acta. 2012; 1819 (644–651).

23. Holwerda SJ, de Laat W. CTCF: the protein, the binding partners, the binding sites and their chromatin loops. Philos Trans R Soc Lond B Biol Sci. 2013; 368: 2012–0369.

24. Caterina JJ, Ryan TM, Pawlik KM, Palmiter RD, Brinster RL, Behringer RR, et al. Human beta-globin locus control region: analysis of the 5’ DNase I hypersensitive site HS 2 in transgenic mice. Proc Natl Acad Sci U S A. 1991 88: 1626–30.

25. Peterson KR, Clegg CH, Navas PA, Norton EJ, Kimbrough TG, Stamatoyannopoulos G. Effect of deletion of 5’HS3 or 5’HS2 of the human beta-globin locus control region on the developmental regulation of globin gene expression in beta-globin locus yeast artificial chromosome transgenic mice. Proc Natl Acad Sci U S A. 1996 93: 6605–9.

26. Navas PA, Peterson KR, Li Q, McArthur M, Stamatoyannopoulos G. The 5’HS4 core element of the human beta-globin locus control region is required for high-level globin gene expression in definitive but not in primitive erythropoiesis. J Mol Biol. 2001 312: 17–26.

27. Farrell CM, West AG, Felsenfeld G. Conserved CTCF insulator elements flank the mouse and human beta-globin loci. Mol Cell Biol. 2002 22: 3820–31.

28. Dostie J, Richmond TA, Arnaout RA, Selzer RR, Lee WL, Honan TA, et al. Chromosome Conformation Capture Carbon Copy (5C): a massively parallel solution for mapping interactions between genomic elements. Genome Res. 2006 16: 1299–309.

29. Chien R, Zeng W, Kawauchi S, Bender MA, Santos R, Gregson HC, et al. Cohesin mediates chromatin interactions that regulate mammalian β-globin expression. J Biol Chem. 2011 286: 17870–8.

30. Wu X, Scott DA, Kriz AJ, Chiu AC, Hsu PD, Dadon DB, et al. Genome-wide binding of the CRISPR endonuclease Cas9 in mammalian cells. Nat Biotechnol. 2014; 32: 670–6. doi: 10.1038/nbt.2889.

31. Kuscu C, Arslan S, Singh R, Thorpe J, Adli M. Genome-wide analysis reveals characteristics of off-target sites bound by the Cas9 endonuclease. Nat Biotechnol. 2014; 32: 677–83. doi: 10.1038/nbt.2916.

32. Cencic R, Miura H, Malina A, Robert F, Ethier S, Schmeing TM, et al. Protospacer adjacent motif (PAM)-distal sequences engage CRISPR Cas9 DNA target cleavage. PLoS One. 2014; 9: e109213. doi: 10.1371/journal.pone.0109213.

33. O’Green H, Henry IM, Bhakta MS, Meckler JF, Segal DJ. A genome-wide analysis of Cas9 binding specificity using ChIP-seq and targeted sequence capture. Nucleic Acids Res. 2015; 43: 3389–404. doi: 10.1093/nar/gkv137.

34. Deng B, Melnik S, Cook PR. Transcription factories, chromatin loops, and the dysregulation of gene expression in malignancy. Semin Cancer Biol. 2013 23: 65–71.

35. Williamson I, Berlivet S, Eskeland R, Boyle S, Illingworth RS, Paquette D, et al. Spatial genome organization: contrasting views from chromosome conformation capture and fluorescence in situ hybridization. Genes Dev. 2014 28: 2778–91.

36. Esvelt KM, Mali P, Braff JL, Moosburner M, Yaung SJ, Church GM. Orthogonal Cas9 proteins for RNA-guided gene regulation and editing. Nat Methods. 2013; 10: 1116–21. doi: 10.1038/nmeth.2681.

37. Mali P, Yang L, Esvelt KM, Aach J, Guell M, DiCarlo JE, et al. RNA-guided human genome engineering via Cas9. Science. 2013; 339: 823–6. doi: 10.1126/science.1232033.

38. Miller AD, Buttimore C. Redesign of retrovirus packaging cell lines to avoid recombination leading to helper virus production. Mol Cell Biol 1986 6: 2895–902.

39. Yamashita R, Sathira NP, Kanai A, Tanimoto K, Arauchi T, Tanaka Y, et al. Genome-wide characterization of transcriptional start sites in humans by integrative transcriptome analysis. Genome Res. 2011 21: 775–89.

40. Seki M, Masaki H, Arauchi T, Makauchi H, Sugano S, Suzuki Y. A comparison of the Rest complex binding patterns in embryonic stem cells and epiblast stem cells. PLoS One. 2014; 9(4): e95374.

41. Fujita T, Ryser S, Tortola S, Piuz I, Schlegel W. Gene-specific recruitment of positive and negative elongation factors during stimulated transcription of the MKP-1 gene in neuroendocrine cells. Nucleic Acids Res. 2007 35: 1007–17.

42. Fujita T, Fujii H. Transcription start sites and usage of the first exon of mouse Foxp3 gene. Mol Biol Rep. 2012 39: 9613–9.

